# Paradoxical improvement of cognitive control in older adults under dual-task walking conditions is associated with more flexible reallocation of neural resources: A Mobile Brain-Body Imaging (MoBI) study

**DOI:** 10.1101/2022.12.14.520469

**Authors:** Eleni Patelaki, John J. Foxe, Emma P. Mantel, George Kassis, Edward G. Freedman

## Abstract

Combining walking with a demanding cognitive task is traditionally expected to elicit decrements in gait and/or cognitive task performance. However, it was recently shown that, in a cohort of young adults, most participants ‘paradoxically’ improved performance when walking was added to performance of a Go/NoGo response inhibition task. The present study aims to extend these previous findings to an older adult cohort, to investigate whether this paradoxical improvement when dual-tasking is observed in healthy older adults. Mobile Brain/Body Imaging (MoBI) was used to record electroencephalographic (EEG) activity, three-dimensional (3D) gait kinematics and behavioral responses in the Go/NoGo task, during sitting or walking on a treadmill, in 34 young adults and 37 older adults. Increased response accuracy during walking, independent of age, was found to correlate with slower responses to stimuli and with walking-related EEG amplitude modulations over latencies and topographies related to the cognitive component of inhibition. On the other hand, aging, independent of response accuracy during walking, was found to correlate with slower treadmill walking speeds and attenuation in walking-related EEG amplitude modulations over latencies and topographies associated with the motor component of inhibition. Older adults whose response accuracy improved during walking manifested neural signatures of both behavioral improvement and aging, suggesting that their flexibility in reallocating neural resources while walking might be maintained for the cognitive but not for the motor inhibitory component. These distinct neural signatures of aging and behavior can potentially be used to identify ‘super-agers’, or individuals at risk for cognitive decline due to aging or neurodegenerative disease.

## Introduction

Aging has been typically associated with loss of cognitive flexibility (1-3), neuronal loss (4-7) and disruption of functional connectivity between brain regions (8, 9). Especially during the earlier stages, modest decline in various executive function domains may be masked by ongoing adaptive processes that compensate for aging-related structural and functional deficits. Employing a paradigm that combines cognitive task performance while walking, a special implementation of dual-tasking (10-12), provides a way to systematically ‘load’ neural circuits and may help to unmask emerging decline in cognitive and gait-motoric domains (13-15). In most cases, exposing younger and older adults to dual-task conditions that combine a walking challenge with a demanding cognitive task has been shown to elicit decrements in gait and/or cognitive task performance in the older compared to the younger group, consistent with the ‘cognitive-motor interference’ (CMI) hypothesis (1, 16-28). There are also studies, however, reporting absence of such dual-task-related deterioration with age under certain dual-task walking conditions (29-31). As such, while dual-task load appears to tax neural resources beyond compensatory capacity in most studies of aging individuals, this is not always the case.

Exceeding compensatory capacity is undoubtedly a function of the complexity of the cognitive task, the intensity and modality of the walking task (e.g. overground versus treadmill walking), the cognitive flexibility and fitness level of each individual, and the age-range of the older group (20, 32-35).

Even within groups of people spanning a narrow age-range, there is variability in behavioral performance with associated differences in neural activity. Previous studies have uncovered associations between distinct behaviors and event-related potential (ERP) amplitude and latency signatures in young adults (36-38). Work from our group showed clear links between behavior and neurophysiology in a dual-task context (39), whereby ‘paradoxical’ improvement in response accuracy during walking in young adults was accompanied by frontal ERP amplitude modulations during key stages of inhibitory processing. Walking-related neural activity changes were absent in young adults whose response accuracy either declined or did not change significantly when dual-tasking (39). Based on previous work in a cohort of older adults, it is anticipated that, at the group level, there will be deterioration in cognitive task performance during walking compared to sitting, aligning with CMI (1, 17, 21). However, it remains to be seen whether all older adults show performance deterioration when walking, or whether, like younger adults, some older adults will actually improve their performance. Our research group recently (39) showed that, in a young adult cohort in which most participants improved performance during walking, there were a few individuals experiencing clear walking-related performance decrements. Here we extend these previous findings to an older, healthy adult cohort. The goal is to determine whether some portion of the older adult population will show cognitive control improvements when walking and to identify objective neural markers of this improvement. Such measures may prove useful as predictive markers of those who have resilience against future cognitive deficits versus those at greater risk for cognitive impairment, and could also shed light on potential adaptive strategies for therapeutic targeting.

As mentioned above, aging has been repeatedly linked to weakening of executive functions (13, 40-42). One of the canonical executive control functions, the ability to inhibit an inappropriate response, is known to be impaired by age-related neurodegenerative processes such as Parkinson’s disease (43-46). Coupling response inhibition tasks with walking has been shown to cause pronounced performance declines and increased competition in prefrontal neural circuits in older adults (47). One commonly used approach to study response inhibition is the so-called visual Go/NoGo task (1, 39, 48-54). These tasks typically involve setting up a response regime whereby participants must respond very regularly to the great majority of stimuli presented (‘Go’ stimuli), such that there is a prepotent inclination to execute such a response, while introducing occasional ‘lure’ NoGo stimuli that require participants to withhold their response. During successful ‘NoGo’ trials where the participant successfully inhibits their response, two prominent stimulus-locked ERP components (known as the N2 and the P3) are observed. The N2 is a negative voltage deflection peaking around 200-350 ms post-stimulus-onset (55-57) and is associated with conflict monitoring processes (52, 54, 58, 59). The N2 has a midline frontocentral scalp distribution and its major generators have been localized to the anterior cingulate cortex (ACC) (54, 55, 58-61). The subsequent P3 component complex is seen as a positive voltage deflection at latencies between 300-600 ms post-stimulus-onset (48, 62), and is characterized by a broad scalp distribution spanning parietal, central and frontal scalp regions (48, 63, 64). There is evidence that during the P3 stage, both motor and cognitive components of inhibitory processing are implemented (63, 65). Apart from reflecting suppression of the prepotent button press, which has been localized to motor and mid-cingulate cortical generators (59, 64, 66-69), the P3 ERP component has also been linked to top-down adjustments in inhibitory behavior subserved by predominantly left-lateralized prefrontal sources (53, 64, 66, 67).

When paired with walking, the visual Go/NoGo response inhibition task has been shown to effectively distinguish between younger and older adults in terms of dual-task-related changes, both in response accuracy and in N2/P3 amplitudes and latencies during successful inhibitions (1). In young adults, on average, preservation or improvement of response accuracy during walking compared to sitting, was accompanied by reduced walking-related N2 and P3 amplitudes (1, 39, 48). On the other hand, older adults exhibited, on average, significant response accuracy reduction during walking, but also attenuated ERP amplitude differences between sitting and walking (1). These findings were interpreted as illustrating an age-related reduction in flexibly adapting neural processes in response to the increased task demands imposed by walking.

Here, we set out to establish whether some minority of older healthy adult participants would show a similar paradoxical improvement in response inhibition task performance during walking to that which we had observed in a significant proportion of younger adults in our prior work. If so, the next obvious question was how such improvers might reallocate neural resources when dual-tasking compared to the older adults who show the more typical performance decrements under the same conditions. A general hypothesis here was that we would observe more flexible reallocation of neural resources, manifested as amplitude modulations in the N2 and P3 components during walking, in those older adults who showed improvement, akin to patterns observed in young adult improvers. Elucidating the neural underpinnings of this dual-task-related improvement can provide a deeper insight into how healthy aging impacts the reallocation of neural resources when task demands increase, and why some people seem to be able to maintain flexibility in reallocating resources across age whereas others cannot.

## Materials and Methods

### Participants

Thirty-four (34) young adults (18-30 years old; age = 22.09 ± 3.12 years; 17 female, 17 male; 30 right-handed, 4 left-handed) and thirty-seven (37) older adults (62-79 years old; age = 70.32 ± 4.54 years; 16 female, 21 male; 29 right-handed, 8 left-handed) participated in the study. Twenty-six (26) of the 34 young adults were common between the present study and Patelaki and colleagues (39). The Montreal Cognitive Assessment (MoCA) was administered to older adults, to ensure that no aging-related cognitive impairment was present. All individuals included in the older cohort scored ≥20 in the MoCA, since a cut-off of 20 has been shown to maximize diagnostic accuracy (70). One 70-year-old adult was excluded from the older cohort due to a MoCA score of 17 (this individual was not counted as a member of the 37-patricipant older cohort). On average, adults of the older cohort scored 26.41 ± 2.20 on the MoCA (score range = 20-30, maximum possible MoCA score = 30).

All participants provided written informed consent, reported no diagnosed neurological disorders, no recent head injuries, and normal or corrected-to-normal vision. The Institutional Review Board of the University of Rochester approved the experimental procedures (STUDY00001952). All procedures were compliant with the principles laid out in the Declaration of Helsinki for the responsible conduct of research. Participants were paid the lab-standard hourly rate for time spent in the lab.

### Experimental Design

A Go/NoGo response inhibition cognitive task was employed. During each experimental block, images were presented in the central visual field for 67 ms with a fixed stimulus-onset-asynchrony of 1017 ms. On average, images subtended 20° horizontally by 16° vertically. The task was coded using the Presentation software (version 20.1, Neurobehavioral Systems, Albany, CA, USA). Participants were instructed to press the button of a wireless game controller using their dominant hand as fast and accurately as possible if the presented image was different from the preceding image (‘Go’ trial). They were instructed to withhold pressing the button if the presented image was the same as the preceding image (‘NoGo’ trial) (Fig. 1). Participants performed blocks of 240 trials in which 209 (87%) were Go trials and 31 (13%) were NoGo trials. NoGo trials were randomly distributed within each block, and it was ensured that NoGo pictures were never presented consecutively.

**Fig. 1.**
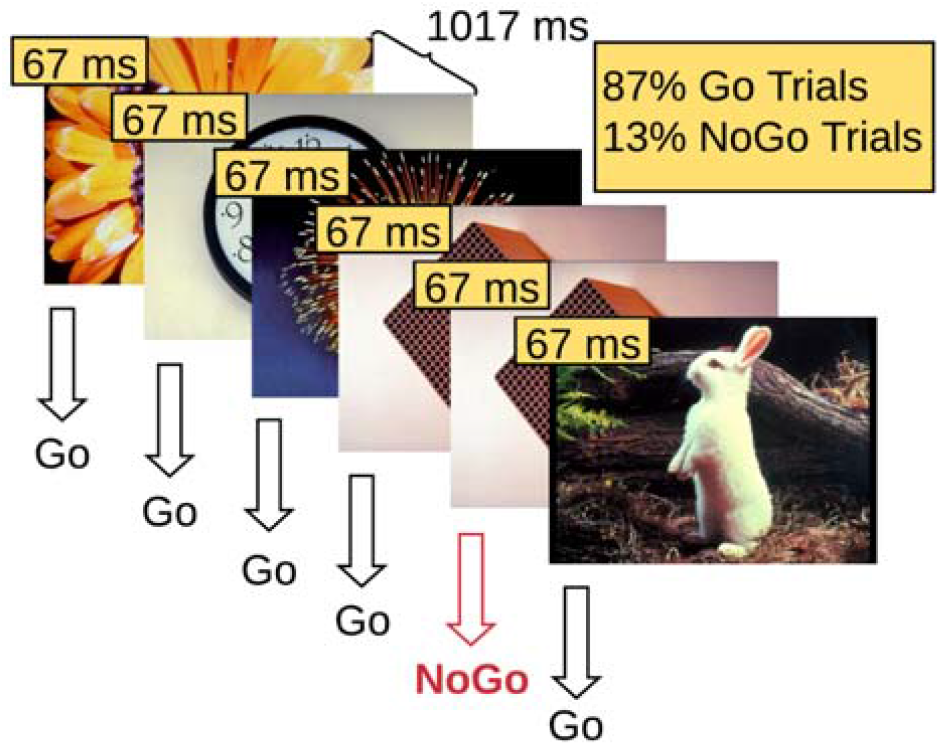
Illustration of the Go/NoGo response inhibition experimental design. Participants are instructed to respond on Go trials and withhold response on NoGo trials.

Four behavioral conditions of the task were defined: 1) correct rejections, defined as the NoGo trials on which participants successfully withheld their response, 2) false alarms, defined as the NoGo trials on which participants incorrectly pressed the response button, 3) hits, defined as the Go trials on which participants correctly pressed the response button, and 4) misses, defined as the Go trials on which participants failed to press the response button.

Experimental blocks were performed while the participants were either sitting or walking on a treadmill (Tuff Tread, Conroe, TX, USA), at a distance of 2.25 m approximately from the projection screen on which the images were projected (Barco F35 AS3D, 1920×1080 pixels). A safety harness was worn while walking to guard against falls (https://youtu.be/HS-5Qk5tvDE). An experimental session consisted of 16 blocks: 1 training block at the beginning, 7 sitting blocks, 7 walking blocks, and a walking-only block (walking on the treadmill without a cognitive task). The order of sitting and walking blocks was pseudorandomized; no more than 3 consecutive walking blocks occurred to prevent fatiguing the participants. Participants were allowed to take short breaks between the blocks. Blocks lasted 4 minutes each. Most participants took at least 1 break during the experiment. If a break was requested, typically it did not last longer than 10 minutes. Participants were asked to select a treadmill speed corresponding to brisk walking for them. On average, young adults walked at 4.64 ± 0.42 km/h, and older adults at 2.74 ± 0.84 km/h.

The pictures used for stimuli were drawn from the International Affective Picture System (IAPS) database (71). The IAPS database contains pictures of varied emotional valence and semantic content. Positive, neutral and negative pictures were all used, however analyzing the emotional valence or semantic content of stimuli is beyond the scope of this study.

EEG data were recorded using a BioSemi Active Two System (BioSemi Inc., Amsterdam, The Netherlands) and a 64-electrode configuration following the International 10-20 system. Neural activity was digitized at 2048 Hz. Full-body motion capture was recorded using a 16 camera OptiTrack system (Prime 41 cameras), and Motive software (OptiTrack, NaturalPoint, Inc., Corvallis, OR, USA) in a ∼37 m^2^ space. Cameras recorded 41 markers on standard anatomical landmarks along the torso, the head and both arms, hands, legs and feet at 360 frames per second. Stimulus triggers from Presentation (Neurobehavioral Systems Inc., Berkeley, CA, USA), behavioral responses from the game controller button, motion tracking data and EEG data were time-synchronized using Lab Streaming Layer (LSL) software (Swartz Center for Computational Neuroscience, University of California, San Diego, CA, USA; available at: https://github.com/sccn/labstreaminglayer). Motion capture data were recorded using custom software written to rebroadcast the data from the Motive software to the LSL lab recorder. EEG data were recorded from available LSL streaming plugins for the BioSemi system. Behavioral event markers were recorded using the built-in LSL functionality in the Presentation software. The long-term test-retest reliability of the MoBI approach has been previously detailed (72). All behavioral, EEG and motion kinematic data processing and basic analyses were performed using custom MATLAB scripts (MathWorks Inc., Natick, MA, USA) and/or functions from EEGLAB (73). Custom analysis code will be made available on GitHub (https://github.com/CNL-R) upon publication.

### Cognitive Task Performance Processing & Analysis

The timing of each button press relative to stimulus onset, the participant’s response times (RTs), were recorded using the Response Manager functionality of Presentation and stored with precision of 1/10 millisecond. The Response Manager was set to accept responses only after 183 ms post-stimulus-onset within each experimental trial. Any responses prior to that were considered delayed responses to the previous trial and were ignored. This RT threshold was selected to filter out as many delayed-response trials as possible, without rejecting any valid trials for which the responses were merely fast (74).

Two behavioral conditions of the cognitive task were examined in terms of EEG activity in this study, 1) correct rejections and 2) hits. For correct rejections, only trials that were preceded by a hit were included to ensure that the inhibitory component was present.

Two behavioral measures were calculated: 1) the d’ score (sensitivity index) and 2) mean RT during (correct) Go trials, namely hits. D’ is a standardized score and it is computed as the difference between the Gaussian standard scores for the false alarm rate (percentage of unsuccessful NoGo trials) and the hit rate (percentage of successful Go trials) (75, 76). D’ was preferred over correct rejection rate (percentage of successful NoGo trials) as a measure of accuracy of inhibitory performance, since it removes the bias introduced by different response strategies adopted across participants. For a more detailed explanation, the reader is referred to the ‘Cognitive task performance processing & analysis’ section of Methods, in Patelaki and colleagues (39).

### EEG Activity Processing & Analysis

EEG signals were first filtered using a zero-phase Chebyshev Type II filter (*filtfilt* function in MATLAB, passband ripple *Apass* = 1 dB, stopband attenuation *Astop* = 65 dB) (77), and subsequently down-sampled from 2048 Hz to 512 Hz. Next, ‘bad’ electrodes were detected based on kurtosis, probability, and spectrum of the recorded data, setting the threshold to 5 standard deviations of the mean, as well as covariance, with the threshold set to ±3 standard deviations of the mean (77). These ‘bad’ electrodes were removed and interpolated based on neighboring electrodes, using spherical interpolation. All the electrodes were re-referenced offline to a common average reference.

It has been shown that 1-2 Hz highpass filtered EEG data yield the optimal Independent Component Analysis (ICA) decomposition results in terms of signal-to-noise ratio (78, 79). In order to both achieve a high-quality ICA decomposition and retain as much low-frequency (< 1 Hz) neural activity as possible, after running Infomax ICA (*runica* function in EEGLAB, the number of retained principal components matched the rank of the EEG data) on 1-45 Hz bandpass-filtered data and obtaining the decomposition matrices (weight and sphere matrices), these matrices were transferred and applied to 0.01-45 Hz bandpass-filtered data. High-pass filtering was selected to be conservative based on evidence that high-pass filters ≤ 0.1 Hz introduce fewer artifacts into the ERP waveforms (80). ICs were labeled using the ICLabel algorithm (81). ICs whose sum of probabilities for the 5 artifactual IC classes (‘Muscle’, ‘Eye’, ‘Heart’, ‘Line Noise’, ‘Channel Noise’) was higher than 50% were labeled as artifacts and were thus rejected. The remaining ICs were back-projected to the sensor space (79, 82).

Subsequently, the resulting neural activity was split into temporal epochs. For both correct rejection and hit trials, epochs were locked to the stimulus onset, beginning 200 ms before and extending until 800 ms after stimulus onset of the trial. Both correct rejection and hit epochs were baseline-corrected relative to the pre-stimulus-onset interval from -100 to 0 ms. Epochs with a maximum voltage greater than ±150 μV or that exceeded 5 standard deviations of the mean in terms of kurtosis and probability were excluded from further analysis. Epochs that deviated from the mean by ±50 dB in the 0-2 Hz frequency window (eye movement detection) and by +25 or -100 dB in the 20-40 Hz frequency window (muscle activity detection) were rejected as well. For the sitting condition, on average 26% of the trials (27% for young adults and 26% for older adults) were rejected based on these criteria, resulting in 940 ± 214 accepted trials for hits (958 ± 191 for young adults and 924 ± 234 for older adults) and 87 ± 37 accepted trials for correct rejections (89 ± 39 for young adults and 85 ± 35 for older adults). For the walking condition the respective percentage was 40% (44% for young adults and 37% for older adults), resulting in 767 ± 272 accepted trials for hits (743 ± 299 for young adults and 788 ± 247 for older adults) and 73 ± 36 accepted trials for correct rejections (74 ± 40 for young adults and 72 ± 32 for older adults). Event-related potentials (ERPs) were measured by averaging epochs for (2 motor-task)x(2 cognitive-task) conditions, namely 4 experimental conditions in total. The motor task conditions were 1) sitting and 2) walking; and the cognitive task conditions were 1) correct rejections and 2) hits.

### Gait Processing & Analysis

Heel markers on each foot were used to track gait kinematics. The three dimensions (3D) of movement were defined as follows: X is the dimension of lateral movement (right-and-left relative to the motion of the treadmill belt), Y is the dimension of vertical movement (up- and-down relative to the motion of the treadmill belt), and Z is the dimension of fore-aft movement (parallel to the motion of the treadmill belt). The heel marker motion in 3D is described by the 3 time series of the marker position over time in the X, Y and Z dimension, respectively. Gait cycle was defined as the time interval between two consecutive heel strikes of the same foot. Heel strikes were identified as the local maxima of the Z position waveform over time. To ensure that no ‘phantom’ heel strikes were captured, only peaks with a prominence greater than 0.1 m were kept (*findpeaks* function in MATLAB, *minimum peak prominence* parameter was set to 0.1 m).

Stride-to-stride variability was quantified as the mean Euclidean distance between consecutive 3D gait cycle trajectories, using the Dynamic Time Warping algorithm (DTW) (83, 84). DTW is an algorithm for measuring the similarity between time series, and its efficacy in measuring 3D gait trajectory similarity is well-established (85-87).

In the case of one-dimensional signals, if Xm=1,2,..,M the reference signal and Yn=1,2,..,N the test signal, then DTW finds a sequence {ix, iy} of indices (called warping path), such that X(ix) and Y(iy) have the smallest possible distance. The ix and iy are monotonically increasing indices to the elements of signals X, Y respectively, such that elements of these signals can be indexed repeatedly as many times as necessary to expand appropriate portions of the signals and thus achieve the optimal match. This concept can be generalized to multidimensional signals, like the 3D gait cycle trajectories of interest here. The minimal distance between the reference and the test signals (gait trajectories here) is given by formula (1):

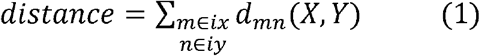

Gait cycle trajectories with a kurtosis that exceeded 5 standard deviations of the mean were rejected as outliers. Also, before DTW computation, gait cycle trajectories were resampled to 100 samples. Since DTW essentially calculates the sum of the Euclidean distances between corresponding points of two interrogated trajectories, ensuring that all trajectories are resampled to the same length helps avoid bias in the algorithm computations.

The actual measure that was used to quantify each participant’s stride-to-stride variability is the mean across DTW distances occurring from all stride-to-stride comparisons. Right-foot and left-foot stride-to-stride DTW distances were pooled to calculate the mean DTW distance per participant.

### Statistical Analyses

#### Cognitive Task Performance

##### Response Accuracy

Walking-*minus*-sitting d’ scores and sitting d’ scores were correlated with age using a Spearman rank correlation, to test potential associations between age and dual-task-related change in response accuracy, and between age and response accuracy during the ‘baseline’ sitting condition, respectively. Due to the bimodal distribution of the age variable, with one of the modes corresponding to young adults (YAs, age range = 18-30 years) and the second to older adults (OAs, age range = 62-79 years), all correlations performed in the context of this study used the non-parametric Spearman rank correlation, since the normality assumption of Pearson’s correlation was violated. If walking-*minus*-sitting d’ scores were found to significantly correlate with age, then d’ score difference between sitting and walking was tested within both age groups (YAs, OAs) using paired t-tests in case of normally distributed data, and Wilcoxon signed rank tests in case of non-normally distributed data. Follow-up testing aimed to determine whether the potential significant correlation was driven by a specific age group.

To allow for a closer inspection of walking-related effects in response accuracy, participants were subsequently classified into 3 groups, based on whether their d’ score during walking was 1) significantly higher than during sitting (d’walking > d’sitting; response accuracy improved significantly while walking; **IMP**), 2) not statistically different from their d’ score during sitting (d’walking ≈ d’sitting; response accuracy did not change significantly while walking; **NC** = No Change), or 3) significantly lower than during sitting (d’walking < d’sitting; response accuracy declined significantly while walking; **DEC**). According to the methodology described in Patelaki and colleagues (39), the individual walking-*minus*-sitting d’ score of each participant was defined as significant if it lay outside of the 95% confidence interval of the normal distribution that had a mean value of zero and a standard deviation equal to that of the walking-*minus*-sitting d’ score distribution of the entire cohort, pooled across age.

##### Response Speed

Walking-*minus*-sitting mean response time (RT) during hits was subjected to 2 partial Spearman rank correlations: 1) with walking-*minus*-sitting d’ score controlling for age, to assess potential associations between dual-task-related response accuracy change and dual-task-related response speed change, free from any effects of age, and 2) with age controlling for walking-minus-sitting d’ score, to test potential associations between age and dual-task-related response speed change, free from any effects of dual-task-related response accuracy change. If either of the correlations above was found to be significant, then mean RT difference between sitting and walking was tested as a follow-up, either within each of the 3 behavioral groups (IMPs, NCs, DECs) if correlation 1 was significant, or within both age groups (YAs, OAs) if correlation 2 was significant. For follow-up testing purposes, paired t-tests were used if the data were normally distributed, and Wilcoxon signed rank tests if the data were non-normally distributed.

#### EEG Activity

The EEG statistical analyses were performed using the FieldTrip toolbox (88) (http://fieldtriptoolbox.org). As mentioned in the Introduction, significant age-related changes in ERP amplitudes during walking were expected to be detected during N2 and P3 latencies during correct rejections (approximately [200, 350] ms and [350,600] ms, respectively (1, 48, 55-57, 62)). However, Malcolm and colleagues (1), who reported these findings, studied these effects only at 3 midline electrode sites, namely FCz (frontocentral), Cz (central) and CPz (centroparietal) electrodes. Conducting statistical analyses only at predetermined latency intervals and a few electrode locations cannot fully elucidate the spatiotemporal distribution of the interrogated effects, therefore the alternative approach of cluster-based permutation tests was employed here (89). Using the same approach, Patelaki and colleagues (39) found significant walking-related ERP amplitude changes in young adults whose response accuracy improved while walking. In that study, cluster-based permutation statistics revealed such walking-related ERP effects during correct rejections over scalp regions that were expected based on the study hypothesis, such as frontocentral regions during N2 latencies, but also over scalp regions that were unanticipated, such as left prefrontal regions during P3 latencies, thereby satisfying both the hypothesis-driven and the exploratory component under a single analysis. The aforementioned walking-related neural activity findings were used for hypothesis generation in the present study, for the arm that aimed to investigate neural correlates of dual-task-related response accuracy changes.

Walking-*minus*-sitting mean ERP amplitudes during hits and correct rejections were calculated at each electrode and timepoint for each participant, by subtracting the within-subject mean sitting ERP waveform (across trials) from the within-subject mean walking ERP waveform. Subsequently, they were subjected to 2 partial Spearman rank correlations at each electrode-timepoint pair: 1) with walking-*minus*-sitting d’ score controlling for age, to assess potential associations between dual-task-related ERP amplitude change and dual-task-related response accuracy change, independent from any effects of age, and 2) with age controlling for walking-minus-sitting d’ score, to test potential associations between dual-task-related ERP amplitude change and age, independent from any effects of dual-task-related response accuracy change. For each of the partial correlations, to identify spatiotemporal clusters of significant neural activity effects while accounting for multiplicity of pointwise electrode-timepoint testing, cluster-based permutation tests were performed using the Monte Carlo method (5000 permutations, significance level of the permutation tests a ::= 0.050, probabilities corrected for performing 2-sided tests) and the weighted cluster mass statistic (90) (cluster significance level a ::= 0.05, non-parametric cluster threshold). The results of the point-wise correlations from all 64 electrodes and all timepoints were displayed as a statistical clusterplot, which is a compact and easily interpretable visualization of the intensity, latency onset/offset, and topography of the detected walking-related ERP effects. The x, y, and z axes, respectively, represent time, electrode location, and the t-statistic (indicated by a color value) at each electrode-timepoint pair. The t-statistic value at electrode-timepoint pairs that did not belong to any cluster of significant effects was masked, namely set to zero (depicted as a teal background).

For Spearman correlations, the t-statistic was calculated based on formula (2) (*r* = correlation coefficient, *N* = sample size) (91):

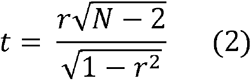

Within either hits or correct rejections, if correlation 1 revealed any clusters of significant effects, then, for each cluster and each of the 3 behavioral groups (IMPs, NCs, DECs), mean ERP amplitude, averaged across electrode-timepoints pairs belonging to the cluster, was follow-up tested for differences between sitting and walking. In a similar manner, within either hits or correct rejections, if correlation 2 revealed any clusters of significant effects, then, for each cluster and each of the age groups (YAs, OAs), mean ERP amplitude, averaged across electrode-timepoints pairs belonging to the corresponding spatiotemporal cluster, was follow-up tested for differences between sitting and walking.

For each correlation that was found to be significant, OA IMPs, which are the group of interest in this study, were follow-up tested for differences between sitting and walking in order to determine whether the effects that they exhibited were characteristic of the age (OA) or behavioral (IMP) group that they belong to. In case correlation 1 was significant, the purpose of the follow-up test would be to assess whether the effects in OA IMPs are consistent with the general IMP effects. In case correlation 2 was significant, the purpose of the follow-up test would be to assess whether the effects in OA IMPs are consistent with the general OA effects.

Follow-up tests were performed using paired t-tests if the data were normally distributed, and Wilcoxon signed rank tests if the data were non-normally distributed.

#### Gait

Two gait-related variables, namely self-selected treadmill walking speed and WT-*minus*-WO mean DTW distance (WT = walking with task, WO = walking only), were subjected to 2 partial Spearman rank correlations each. Each gait-related variable was partially correlated 1) with walking-*minus*-sitting d’ score controlling for age, to assess potential associations between the interrogated gait-related variable and dual-task-related response accuracy change, removing any effects of age, and 2) with age controlling for walking-minus-sitting d’ score, to test potential associations between the interrogated gait-related variable and age, removing any effects of dual-task-related response accuracy change. For WT-*minus*-WO mean DTW distance, if correlation 1 was found to be significant, then mean DTW distance difference between WO and WT was follow-up tested within each of the 3 behavioral groups (IMPs, NCs, DECs); similarly, if correlation 2 was found to be significant, mean DTW distance difference between WO and WT was follow-up tested within both age groups (YAs, OAs). Paired t-tests were used for follow-up testing in case of normally distributed data, and Wilcoxon signed rank tests in case of non-normally distributed data.

It should be pointed out that 1 young and 2 older adults were excluded from the stride-to-stride (DTW) variability analysis due to lack of walking-only kinematic data, thus resulting in a sample size of 68 participants (33 YAs, 35 OAs) for this analysis.

#### Other correlations

To determine whether dual-task-related change in response accuracy could be predicted based on sex, a partial Spearman rank correlation was performed to partially correlate sex with walking-*minus*-sitting d’ score, controlling for age. Furthermore, a Spearman rank correlation was used to correlate MoCA score with walking-*minus*-sitting d’ score, in order to assess whether MoCA scores could predict dual-task-related change in response accuracy.

#### Multiple Comparisons Correction

The 2-stage step-up False Discovery Rate (FDR) method of Benjamini and colleagues (92) was applied on the combined set of all the p-values occurring from all the planned (partial) Spearman rank correlations in cognitive task performance, gait kinematic and EEG activity. The calculations were conducted in MATLAB (93). For each correlation, one p-value was obtained. Specifically, for each of the 4 EEG activity correlation analyses, besides controlling for multiple electrode-timepoint tests within the analysis using cluster-based statistics, the minimum p-value across p-values of all detected clusters was added to the abovementioned combined set for FDR calculation purposes. In total, this combined set comprised of 14 p-values. The significance level yielded by FDR procedure was α_*FDR*_ = 0.0216 (false discovery rate = 5%). Therefore, all correlations whose uncorrected p-values were found to be less than or equal to 0.0216 were significant after FDR correction. Correlations that were found non-significant after FDR correction were not further examined. For those that were found significant, the follow-up paired-sample tests (t-test or Wilcoxon signed rank) that were conducted were not corrected (significance level α = 0.0500).

## Results

### Cognitive Task Performance

#### Response Accuracy

When combining treadmill walking with a Go/NoGo response inhibition task, older adults (OAs) were previously shown to exhibit significantly greater dual-task-related deterioration in response accuracy compared to young adults (YAs), even though their mean response accuracy across motor load condition was found not to differ significantly from that of young adults (1). Here, the same dual task was employed and the potential effect of age on dual-task-related response accuracy change and on ‘baseline’ response accuracy during sitting were tested by correlating participants’ age with walking-minus-sitting d’ scores and with sitting d’ scores, respectively (higher d’ scores indicate better discriminability between Go and NoGo stimuli). While no significant correlation was revealed between age and sitting d’ scores (Spearman’s r = -0.11, p = 0.3625), age was found to negatively correlate with dual-task-related d’ change (Spearman’s r = -0.42, p = 0.0002, α_*FDR*_ = 0.0216) indicating that response accuracy during walking – not response accuracy in the ‘baseline’ sitting condition – deteriorates significantly with age. This is consistent with previous findings (1).

To test whether this significant correlation effect between age and walking-related response accuracy change was driven solely by effects in only one of the two age groups, the impact of walking on response accuracy was assessed within each age group separately using follow-up paired t-tests. In the YA cohort, d’ scores were significantly higher during walking compared to sitting (d’sitting = 2.12 ± 1.27, d’walking = 2.31 ± 1.11; t33 = 2.41, p = 0.0219, α = 0.0500, Cohen’s d = 0.41) indicating that treadmill walking improved young adults’ performance in the Go/NoGo task. In contrast, in the OA cohort, d’ scores were significantly lower during walking (d’sitting = 1.88 ± 0.96, d’walking = 1.66 ± 0.85; t36 = 2.85, p = 0.0023, α = 0.0500, Cohen’s d = 0.54) indicating that older adults’ performance in the Go/NoGo task deteriorated when this was paired with treadmill walking. Based on these findings, walking-related change in response accuracy with age was driven both by improvement in the younger group and by decline in the older group. At a group level, the dual-task-related decline exhibited by OAs is consistent with the ‘cognitive-motor interference’ (CMI) hypothesis, while dual-task-related improvement that YAs exhibit appears to be inconsistent with the CMI hypothesis, though in agreement with previous findings (39).

Fig. 2 shows the distributions of d’ scores during sitting and walking, in each of the two age groups. By inspecting d’ performance at an individual level, it was observed that while most YAs improved when dual-tasking and most OAs declined, there were also participants in each age group whose performance was distinct from the group average. Specifically, from the 34 YAs, 20 improved when walking (IMPs), 6 did not change significantly across motor load condition (NCs) and 8 declined when walking (DECs). Interestingly, the distribution of the 37 OAs across these 3 behavioral groups (IMPs, NCs, DECs) was essentially ‘inverted’ compared to the YA distribution, with 7 IMPs, 10 NCs and 20 DECs in the older cohort (Fig. 2). Of note, to determine whether change in response accuracy within each individual participant was significant or not, the method described in Patelaki and colleagues (39) was used. According to this method, the decision was based on whether the walking-*minus*-sitting d’ score of the participant lay outside of the 95% confidence interval of the normal distribution having a mean value of zero and a standard deviation equal to that of the walking-*minus*-sitting d’ score distribution of the entire cohort pooled across age. If d’walking > d’sitting, namely the participant improved significantly during walking, they were classified into the IMP group. If d’walking ≈ d’sitting, namely the participant did not change significantly between walking and sitting, they were classified into the NC group. If d’walking < d’sitting, namely the participant deteriorated significantly during walking, they were classified into the DEC group.

**Fig. 2.**
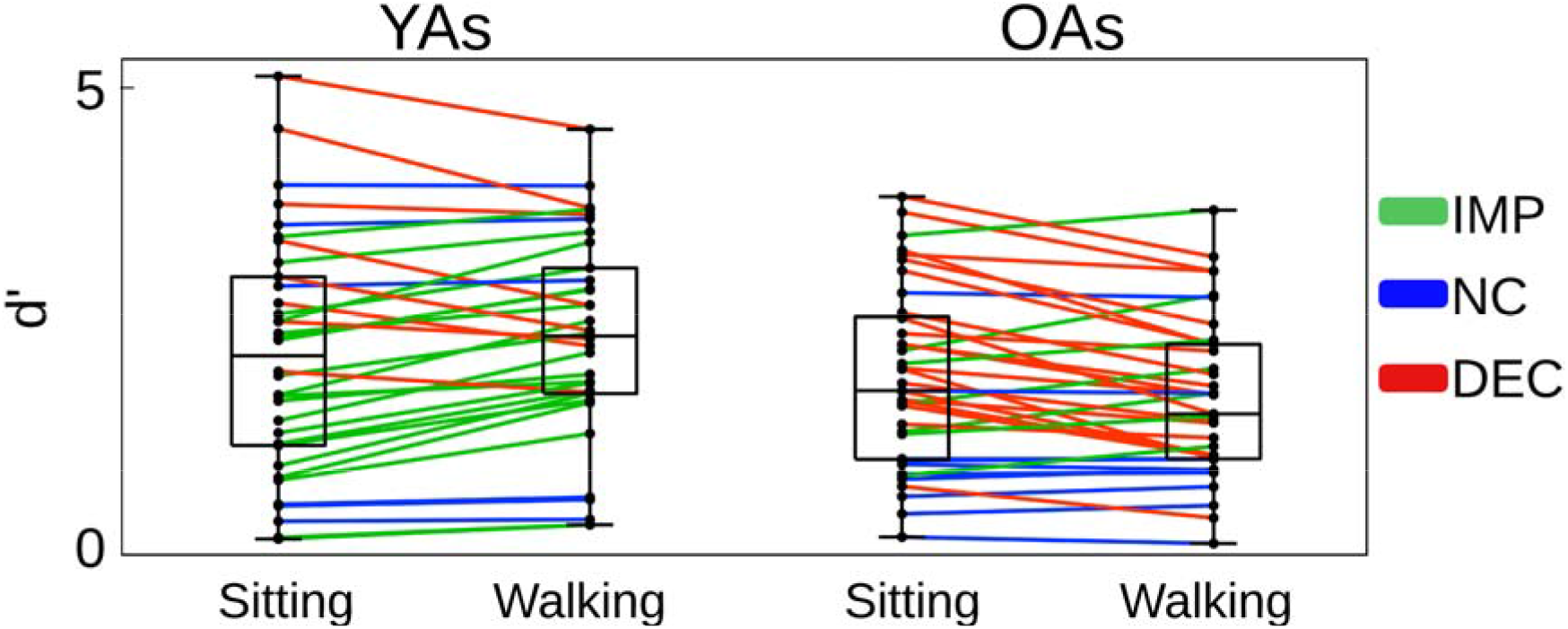
Sitting and walking d’ score distributions, in young adults (YAs) and older adults (OAs). Each line corresponds to one participant. Participants who improved when walking (IMPs; d’walking > d’sitting) are shown in green, those who did not change significantly across motor load condition (NCs; d’walking ≈ d’sitting) are shown in blue, and those who declined when walking (DECs; d’walking > d’sitting) are shown in red. Black dots on the vertical line represent individual participants. The central mark of each box indicates the median, and the bottom and top edges indicate the 25th and 75th percentiles, respectively. The whiskers extend to the most extreme data points not considered outliers. There were no outliers here.

The observation that there were a few (7) older participants whose behavioral response to dual-task load was uncharacteristic of their age, namely they improved during walking, highlighted the need for identifying distinct neural signatures of dual-task-related improvement with aging, to shed light on the neural processes underlying this seemingly paradoxical behavior. To this end, neural signatures of behavioral improvement during dual-task walking, independent of age, were calculated by partially correlating dual-task-related ERP amplitude changes (quantified as walking-*minus*-sitting mean ERP amplitudes) with dual-task-related d’ score changes (quantified as walking-*minus*-sitting d’ scores) while controlling for age. Similarly, to obtain neural signatures of aging, independently of behavior during dual-task walking, dual-task-related ERP activity changes were partially correlated with age while controlling for dual-task-related d’ score changes. ERP activity both during correct rejection trials and during hit trials was tested using these correlation analyses, in order to investigate whether the presence of the inhibitory component (correct rejections), or its absence (hits), played a role in the way that the employed dual task was managed by the neural resources across age. Additionally, to test whether potential changes in other physiological variables besides ERPs are age-related, or they reflect performance trade-offs, the following variables were partially correlated with age and dual-task-related d’ score changes, similarly as above: dual-task-related response speed changes (walking-*minus*-sitting mean response time during hits), dual-task-related stride-to-stride variability changes (quantified as WT-minus-WO mean DTW distance; WT = walking with task, WO = walking only, DTW = Dynamic Time Warping) and self-selected treadmill speed. Finally, to assess whether sex and cognitive assessment (MoCA) scores can predict walking-related change in response accuracy, these 2 outcome variables were correlated with walking-related d’ score changes. Motivated by the consistent finding of Go/NoGo response accuracy decline with age, uniquely observed under walking conditions, most correlation analyses outlined above were focused on dual-task-related effects in each of the interrogated physiological domains, namely gait, neurophysiology, and behavior in the cognitive task.

#### Response Speed

Fig. 3 shows the distributions of mean response times (RT) during sitting and walking, in each of the age groups.

**Fig. 3.**
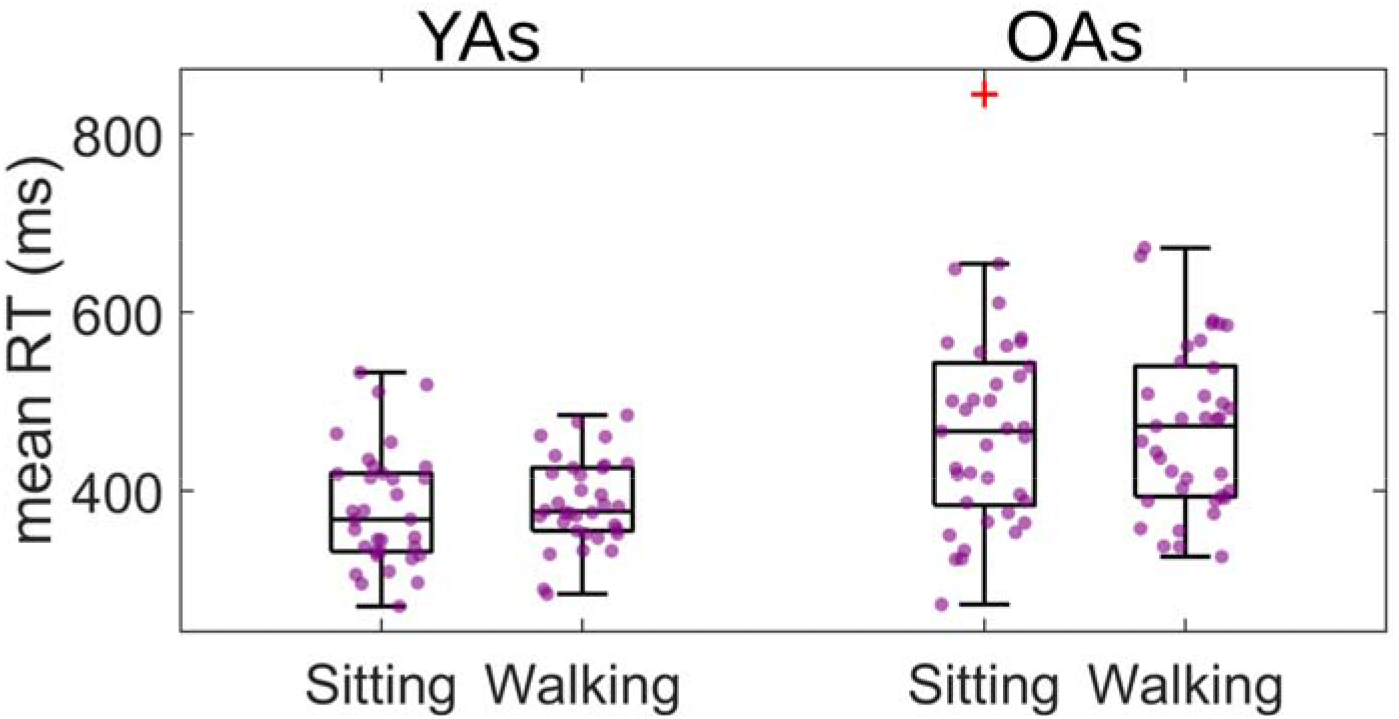
Sitting and walking mean response time (RT) distributions during hits (correct Go trials), in YAs and OAs. Purple dots represent individual participants. Red ‘+’ symbols represent outliers.

Walking-*minus*-sitting mean response time during hits was examined for associations with 1) walking-related change in response accuracy (walking-minus-sitting d’ score), controlling for age, and with 2) age, controlling for walking-related change in response accuracy, using partial Spearman rank correlations. Correlation 1 was found to be significantly positive (Spearman’s r = 0.44, p = 0.0001, α_*FDR*_ = 0.0216), while correlation 2 was found to be non-significant (Spearman’s r = 0.14, p = 0.2493).

To test whether this significant correlation effect between walking-related change in response speed and accuracy was driven by effects in any one of the three behavioral groups (i.e., IMP, NC or DEC), the impact of walking on response speed was assessed within each behavioral group separately using two paired t-tests (IMP and NC groups) and 1 Wilcoxon signed rank test (DEC group). Slower response times were found during walking in IMPs (mean RT sitting = 368 ± 70 ms, mean RT walking = 382 ± 59 ms; t27 = 2.34, p = 0.0273, α = 0.0500, Cohen’s d = 0.45), but no significant differences between sitting and walking were detected either in NCs (p = 0.2098) or DECs (p = 0.3164). This finding indicates that walking-related improvement in response accuracy was accompanied by walking-related slowing in responses to stimuli, suggesting the existence of a speed-accuracy tradeoff when engaging in the dual task.

### EEG Activity

Fig. 4 shows the grand mean ERP waveforms during sitting (blue) and walking (red) at 3 midline electrode locations: FCz, Cz and CPz. Observed ERPs during hits and correct rejections (CRs) are shown for both for younger (A) and for older adults (B). These electrodes were selected for illustration because the N2 and P3 ERP amplitudes during correct rejections are typically found to be maximal over these midline electrode locations (48, 52, 63). All ERP waveforms were aligned on the onset of stimulus presentation (vertical line - time = 0), and they were plotted from 200 ms pre-stimulus-onset to 800 ms post-stimulus-onset. The N2 component is the negative voltage deflection spanning ∼200-350 ms post-stimulus-onset (55-57), and it is evident during both hits and correct rejections. Smaller N2 amplitudes were observed in OAs (Fig. 4B) compared to YAs (Fig. 4A) (1). The P3 component is the positive voltage deflection extending ∼350-600 ms post-stimulus-onset (48, 62), and it was more strongly evoked during correct rejections than during hits in both age groups (1). During correct rejections, in YAs, the P3 component was maximal at Cz and CPz (Fig. 4A). In contrast, OAs exhibited marked P3 deflections at all 3 midline locations (Fig. 4B). This observation suggests anteriorization of the P3 component with aging (1, 94, 95). Subsequent analyses will focus on partially correlating walking-related ERP amplitude changes during hits and correct rejections, first with walking-related d’ score changes while controlling for age, and then with age while controlling for walking-related d’ score changes. This approach permits the dissociation of these two variables in terms of the walking-related neural activity changes that they are linked to.

**Fig. 4.**
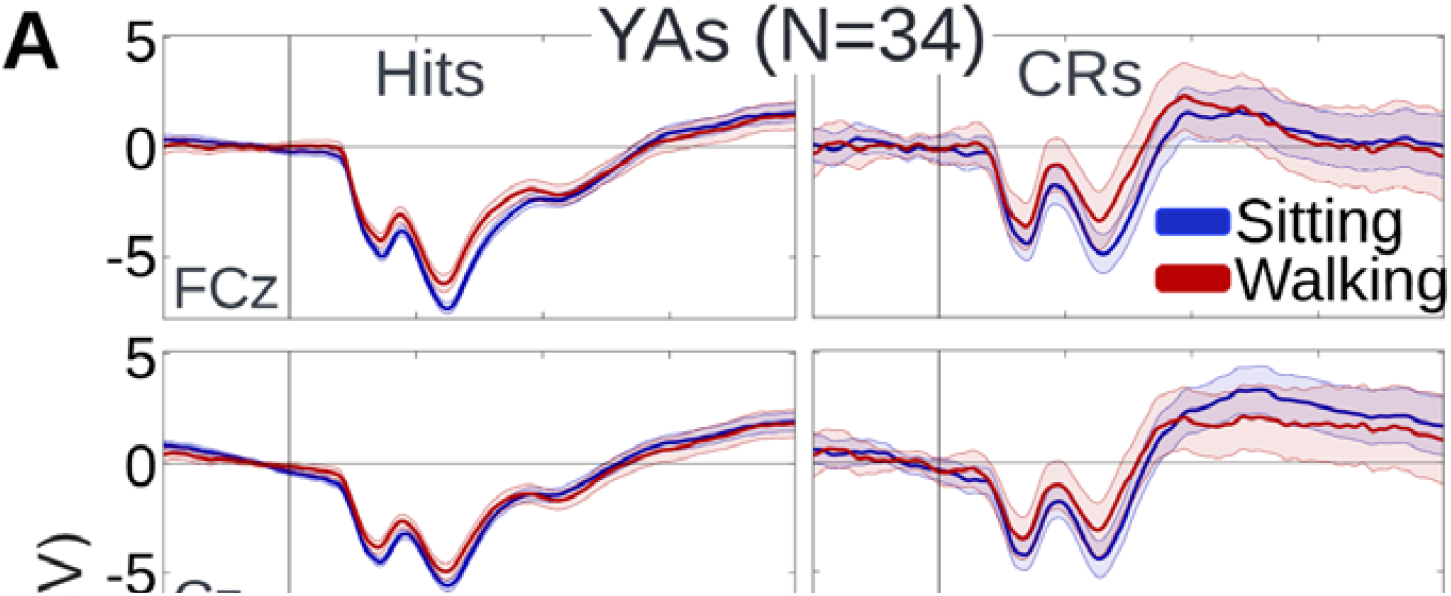
Grand mean sitting (blue) and walking (red) ERP waveforms, at 3 midline electrode locations: frontocentral midline (FCz), central midline (Cz) and centroparietal midline (CPz). ERPs are shown both during hits and during correct rejections (CRs), both for YAs (panel **A**) and for OAs (panel **B**). The shaded regions around the ERP traces indicate the Standard Error of the Mean (SEM) across participants of the same age group.

#### Correct Rejections

##### Partial correlation with walking-minus-sitting d’ score, controlling for age

This section aimed to test whether walking-related changes in neural activity during correct rejections correlated with walking-related change in response accuracy independently of age, and if so, at which scalp regions and during which stages of the inhibitory processing stream.

Walking-*minus*-sitting mean ERP amplitude during correct rejections was calculated at each of the 64 electrodes and each epoch timepoint and it was subsequently partially correlated with walking-minus-sitting d’ score, controlling for age, using partial Spearman rank correlations. Cluster-based permutation tests were used to identify spatiotemporal clusters of significant neural activity effects while accounting for multiple electrode/timepoint comparisons.

During correct rejection trials, walking-*minus*-sitting mean ERP amplitudes were found to positively correlate with walking-*minus*-sitting d’ score over frontal scalp regions, during latencies corresponding to multiple stages of inhibitory processing. These correlation effects are represented by the yellow clusters in the Fig. 5A statistical clusterplot (within-cluster mean Spearman’s r = 0.27, p = 0.0216, α_*FDR*_ = 0.0216). Specifically, significant correlation effects were detected as early as a few milliseconds post-stimulus-onset until the end of the epoch (800 ms post-stimulus-onset). To facilitate the study of these effects, 3 latency intervals were defined based on the existing response inhibition ERP literature: 1) sensory/perceptual interval ([0, 200] ms (96-98)), which encompasses the N1 component, 2) the conflict monitoring interval ([200, 350] ms (55-57)), which includes the N2 component, and 3) the control implementation interval ([350, 800] ms (48, 62)), encompassing the P3 component. In the entire cohort pooled across age, as response accuracy increased during walking, walking-related ERP amplitudes were found to become less negative over frontocentral scalp during intervals 1 and 2, and predominantly left-lateralized prefrontal scalp during interval 3 (Fig. 5A). The latencies corresponding to the 3 intervals are demarcated by black dashed rectangles on the clusterplot.

**Fig. 5.**
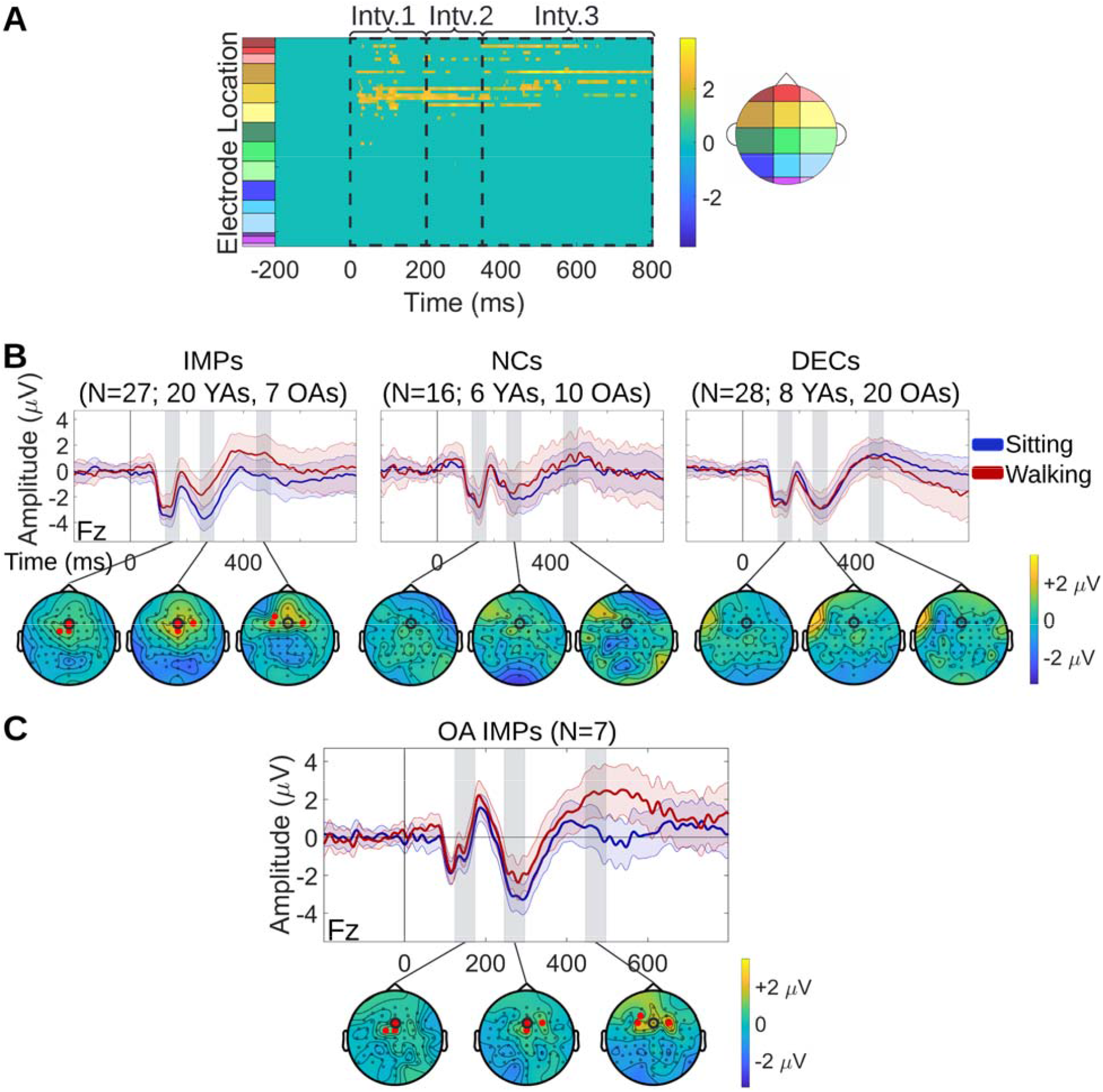
Correlation effects between dual-task-related neural activity change and dual-task-related response accuracy change during correct rejection trials. **A**. Spatiotemporal clusters of walking-*minus*-sitting mean ERP amplitudes correlating with walking-*minus*-sitting d’ score during correct rejections, identified using cluster-based permutation tests. The statistical clusterplot shows the t-values for the electrode-timepoint pairs at which significant correlation was found. Positive t-values (yellow) indicate that, as walking d’ scores increased relative to sitting d’ scores, walking ERP amplitudes became less negative relative to the sitting ERP amplitudes. Such positive correlation effects were found over frontocentral scalp during intervals 1 and 2, and over predominantly left-lateralized prefrontal scalp during interval 3. The black dashed rectangles on the clusterplot demarcate the latencies corresponding to the 3 intervals. **B**. Grand mean sitting and walking ERP waveforms of IMPs (left), NCs (middle) and DECs (right), at a frontal midline electrode (Fz) that exhibited significant correlation effects. The topographical maps show the scalp distribution of the walking-*minus*-sitting mean ERP amplitudes during correct rejections, averaged within each of the behavioral groups (IMPs, NCs, DECs) separately, for 3 selected 50-ms windows (gray bars). Each 50-ms window corresponds to one interval. For each window, the electrodes that exhibited significant correlation effects are indicated by red dots on the topographical map of the behavioral group that was found to drive the correlation. The correlation was driven by IMPs for all 3 intervals. The Fz electrode to which the ERP waveforms correspond is circled in black on the maps. **C**. Follow-up visualizations within the OA IMP group. Grand mean sitting and walking ERP waveforms of OA IMPs at the Fz electrode, along with topographical maps of walking-*minus*-sitting mean ERP amplitudes for the same three 50-ms windows as in panel B.

Follow-up tests were performed to determine whether the effects detected in each of the intervals were driven by a specific type of dual-task-related behavior; in other words, whether the correlation effects were driven by any of the three behavioral groups (IMP, NC, DEC) defined above, in the ‘Cognitive Task Performance’ section. Specifically, for each latency interval and within each behavioral group, the mean ERP amplitude, averaged across electrode-timepoint belonging to the respective significant yellow cluster, was compared between sitting and walking using paired t-tests (or Wilcoxon signed rank tests when the data were non-normally distributed). Such significant ERP amplitude differences between sitting and walking were found in all 3 latency intervals for IMPs (20 YAs, 7 OAs; pInterval-1 = 0.0002, pInterval-2 < 0.0001, pInterval-3 = 0.0008, α = 0.0500), but in none of the 3 intervals for NCs (6 YAs, 10 OAs; pInterval-1 = 0.4321, pInterval-2 = 0.1958, pInterval-3 = 0.8881) or DECs (8 YAs, 20 OAs; pInterval-1 = 0.1791, pInterval-2 = 0.9516, pInterval-3 = 0.6164). These results indicate that correlation effects were driven by significantly reduced ERP amplitudes during intervals 1 and 2, and significantly increased ERP amplitudes during interval 3, in IMPs when walking. The topographical maps of Fig. 5B show the scalp distribution of the walking-*minus*-sitting mean ERP amplitudes during correct rejection trials, averaged across IMPs, NCs and DECs, separately, for three 50-ms windows indicated by gray bars. These 50-ms windows were centered at the peak latency of N1, N2 and P3, respectively, calculated based on the grand mean waveform at FCz, averaged across behavior, age and motor load. N1 peaked at 140 ms, N2 at 270 ms and P3 at 470 ms post-stimulus-onset, approximately. For each window, the electrodes that exhibited significant correlation effects are denoted by red dots on the topographical map of the behavioral group that was found to drive the correlation. The correlation was driven by IMPs for all 3 intervals. Sitting and walking ERP waveforms during correct rejection trials for each behavioral group are depicted at Fz (frontal midline), since this electrode exhibited significant correlation effects (Fig. 5B).

Walking-related ERP amplitude changes in OA IMPs were follow-up tested in the clusters of each of the intervals, to assess how the effects in this group compare to those in the general IMP group. For each latency interval, the mean ERP amplitude was compared between sitting and walking. Significant differences were found in all 3 latency intervals (pInterval-1 = 0.0169, pInterval-2 = 0.0294, pInterval-3 = 0.0081, α = 0.0500), consistent with the findings in the general IMP group. Fig. 5C illustrates the neural activity effects exhibited by OA IMPs. The same ERP waveforms and topographical maps as in Fig. 5B are displayed for OA IMPs only. Indeed, the positive walking-*minus*-sitting mean ERP amplitudes reflecting less negative ERP amplitudes during walking, shown in yellow on the topographical maps, were present over cluster-related frontal topographies in OA IMPs.

These data indicate that reduced walking-related ERP amplitudes over frontocentral regions during the N1and N2 stages of correct rejections, and increased walking-related ERP amplitudes over predominantly left-lateralized prefrontal regions during the P3 stage, constitute neural signatures of behavioral improvement in the cognitive task, independently of age.

##### Partial correlation with age, controlling for walking-minus-sitting d’ score

Walking-related changes in neural activity during correct rejections were tested for correlations with age, independently of walking-related change in response accuracy, across the entire 64-electode set and all the epoch timepoints. Walking-*minus*-sitting mean ERP amplitude during correct rejections was partially correlated with age at each electrode-timepoint pair, controlling for walking-minus-sitting d’ score, using partial Spearman rank correlations. Cluster-based permutation tests were used to identify spatiotemporal clusters of significant neural activity effects while accounting for multiple electrode/timepoint comparisons. For consistency purposes, the same interval definitions as in the previous section are used here too.

During correct rejection trials, walking-*minus*-sitting mean ERP amplitudes were found to negatively correlate with age over predominantly left-lateralized frontal scalp regions during interval 2 and early interval 3. These negative correlation effects are represented by the blue cluster in the Fig. 6A statistical clusterplot (within-cluster mean Spearman’s r = -0.23, p = 0.0120, α_*FDR*_ = 0.0216). Of note, the portion of this blue cluster which extended into interval 3 was excluded from further analysis, because it was considered to add too little value due to its short duration and absence of any novel effects relative the interval-2 portion.

**Fig. 6.**
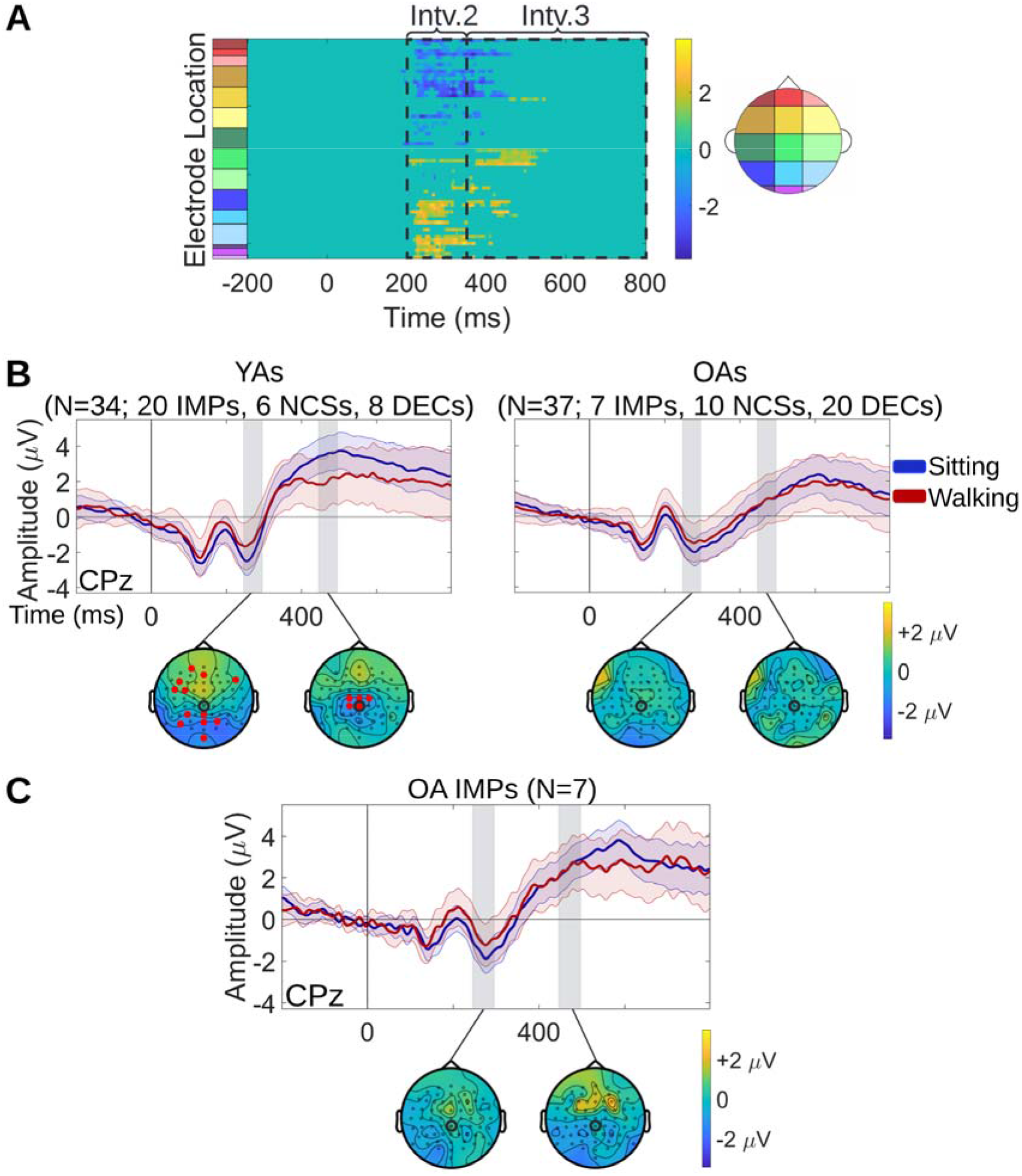
Correlation effects between dual-task-related neural activity change and age during correct rejection trials. **A**. Spatiotemporal clusters of walking-*minus*-sitting mean ERP amplitudes correlating with age during correct rejections, identified using cluster-based permutation tests. The statistical clusterplot shows the t-values for the electrode-timepoint pairs at which significant correlation was found. Negative t-values (blue) indicate that, as age increased, walking ERP amplitudes became more negative relative to the sitting ERP amplitudes. Negative correlation effects were found over predominantly left-lateralized frontal scalp regions during interval 2 and early interval 3. Positive t-values (yellow) indicate that, as age increased, walking ERP amplitudes became more positive relative to the sitting ERP amplitudes. Positive correlation effects were found over parietooccipital regions during interval 2 and over centroparietal regions during interval 3. The latencies of intervals 2 and 3 are demarcated by black dashed rectangles on the clusterplot. **B**. Grand mean sitting and walking ERP waveforms of YAs (left) and OAs (right), at CPz that exhibited significant correlation effects. The topographical maps show the scalp distribution of the walking-*minus*-sitting mean ERP amplitudes during correct rejections, averaged within each of the age groups (YAs, OAs) separately, for two 50-ms window (gray bars). These 2 windows correspond to intervals 2 and 3, respectively, and they are the same as in Fig. 5. For each window, the electrodes that exhibited significant correlation effects are indicated by red dots on the topographical map of the age group that was found to drive the correlation. The correlation was driven by YAs for both intervals. The CPz electrode to which the ERP waveforms correspond is circled in black on the maps. **C**. Follow-up visualizations for the OA IMP group. Grand mean sitting and walking ERP waveforms of OA IMPs at the CPz electrode, along with topographical maps of walking-*minus*-sitting mean ERP amplitudes for the same 50-ms windows as in panel B.

Additionally, positive correlations with age were found over parietooccipital regions during interval 2 and over centroparietal regions during interval 3. These positive correlation effects are represented by the yellow cluster in the Fig. 6A statistical clusterplot (within-cluster mean Spearman’s r = 0.22, p = 0.0160, α_*FDR*_ = 0.0216). All correlation effects detected here reflect age-related increase in ERP amplitudes during walking relative to sitting. Specifically, at electrodes and latencies where ERP amplitudes were negative, they became more negative with age during walking relative to sitting; similarly, at electrodes and latencies where ERP amplitudes were positive, they became more positive with age during walking relative to sitting. The latencies corresponding to intervals 2 and 3 are demarcated by black dashed rectangles on the clusterplot.

Follow-up tests were conducted to determine whether the detected effects were driven by either of the YA or OA age group. For YAs and OAs separately, the mean ERP amplitude, averaged across electrode-timepoint pairs, was compared between sitting and walking within the significant blue cluster of interval 2 and within the significant yellow clusters of intervals 2 and 3 using paired t-tests (or Wilcoxon signed rank tests when the data were non-normally distributed). Significant amplitude differences were found in YAs (20 IMPs, 6 NCs, 8 DECs; pInterval-2-blue = 0.0004, pInterval-2-yellow = 0.0001, pInterval-3-yellow < 0.0001, α = 0.0500), but not for OAs (7 IMPs, 10 NCs, 20 DECs; p_Interval-2-blue_ = 0.7954, p_Interval-2-yellow_ = 0.2675, p_Interval-3-yellow_ = 0.5614). Based on these results, correlation effects were driven by significantly reduced ERP amplitudes in YAs when walking, during both intervals 2 and 3. The topographical maps of Fig. 6B show the scalp distribution of the walking-*minus*-sitting mean ERP amplitudes during correct rejection trials, averaged across YAs and OAs, separately, for 2 selected 50-ms windows (gray bars). These windows correspond to intervals 2 and 3, and they are centered at 270 ms and 470 ms post-stimulus-onset, respectively, the same convention as used in Fig. 5. The electrodes that exhibited significant correlation effects within each of the 2 windows are denoted by red dots on the topographical map of the age group that was found to drive the correlation, which is YAs for both intervals. Sitting and walking ERP waveforms during correct rejection trials for each age group are depicted at CPz, since this electrode exhibited significant correlation effects (Fig. 6B). Walking-related ERP amplitude changes in OA IMPs were follow-up tested in the clusters of each of the 2 intervals, to assess how the effects in this group compare to those in the general OA group. For each latency interval, the mean ERP amplitude was compared between sitting and walking. No detectable differences were found in either interval (pInterval-2-blue = 0.9695, pInterval-2-yellow = 0.5965, pInterval-3-yellow = 0.6029), consistent with the findings in the general OA group. Fig. 6C illustrates the neural activity effects exhibited by OA IMPs. The same ERP waveforms and topographical maps as in Fig. 6B are displayed for OA IMPs only. Indeed, the walking-*minus*-sitting mean ERP amplitudes appeared attenuated in OA IMPs over cluster-related scalp topographies.

Based on these data, attenuations of walking-related ERP amplitude modulations over parietooccipital and predominantly left-lateralized frontal regions during the N2 stage, and over centroparietal regions during the P3 stage are revealed as neural signatures of aging, independently of walking-related behavioral change in the cognitive task.

#### Hits

##### Partial correlation with walking-minus-sitting d’ score, controlling for age

Walking-related changes in neural activity during hits were tested for correlations with walking-related change in response accuracy independently of age, across the entire 64-electode set and all the epoch timepoints. Walking-*minus*-sitting mean ERP amplitude during hits was calculated at each electrode-timepoint pair and it was subsequently partially correlated with walking-minus-sitting d’ score, controlling for age, using partial Spearman rank correlations. Cluster-based permutation tests revealed no clusters of significant spatiotemporal neural activity effects.

##### Partial correlation with age, controlling for walking-minus-sitting d’ score

Walking-related changes in neural activity during hits were tested for correlations with age independently of walking-related change in response accuracy, across the entire 64-electode set and all the epoch timepoints. Walking-*minus*-sitting mean ERP amplitude during hits was partially correlated with age at each electrode-timepoint pair, controlling for walking-minus-sitting d’ score, using partial Spearman rank correlations. Cluster-based permutation tests revealed one cluster of significant spatiotemporal neural activity effects spanning left-lateralized centroparietal regions, which, however, did not remain significant after correction for multiple comparisons (p = 0.0468, α_*FDR*_ = 0.0216).

### Gait

#### Treadmill Walking Speed

Self-selected treadmill walking speed was tested for associations with 1) walking-minus-sitting d’ score, controlling for age, and with 2) age, controlling for walking-minus-sitting d’ score, using partial Spearman rank correlations. While correlation 1 was found to be non-significant (Spearman’s r = -0.02, p = 0.8684), correlation 2 was found to be significantly negative (Spearman’s r = -0.68, p < 0.0001, α_*FDR*_ = 0.0216), indicating that selection of slower walking speeds on the treadmill is driven by older age and not behavioral performance in the dual task.

#### Stride-to-Stride Variability

Fig. 7 demonstrates the stride-to-stride variability distributions of young and older adults, with and without a concurrent cognitive task (WT and WO, respectively) (panel B), along with the 3D representations of trajectories of a series of strides for 1 young and 2 older participants (panel A).

**Fig. 7.**
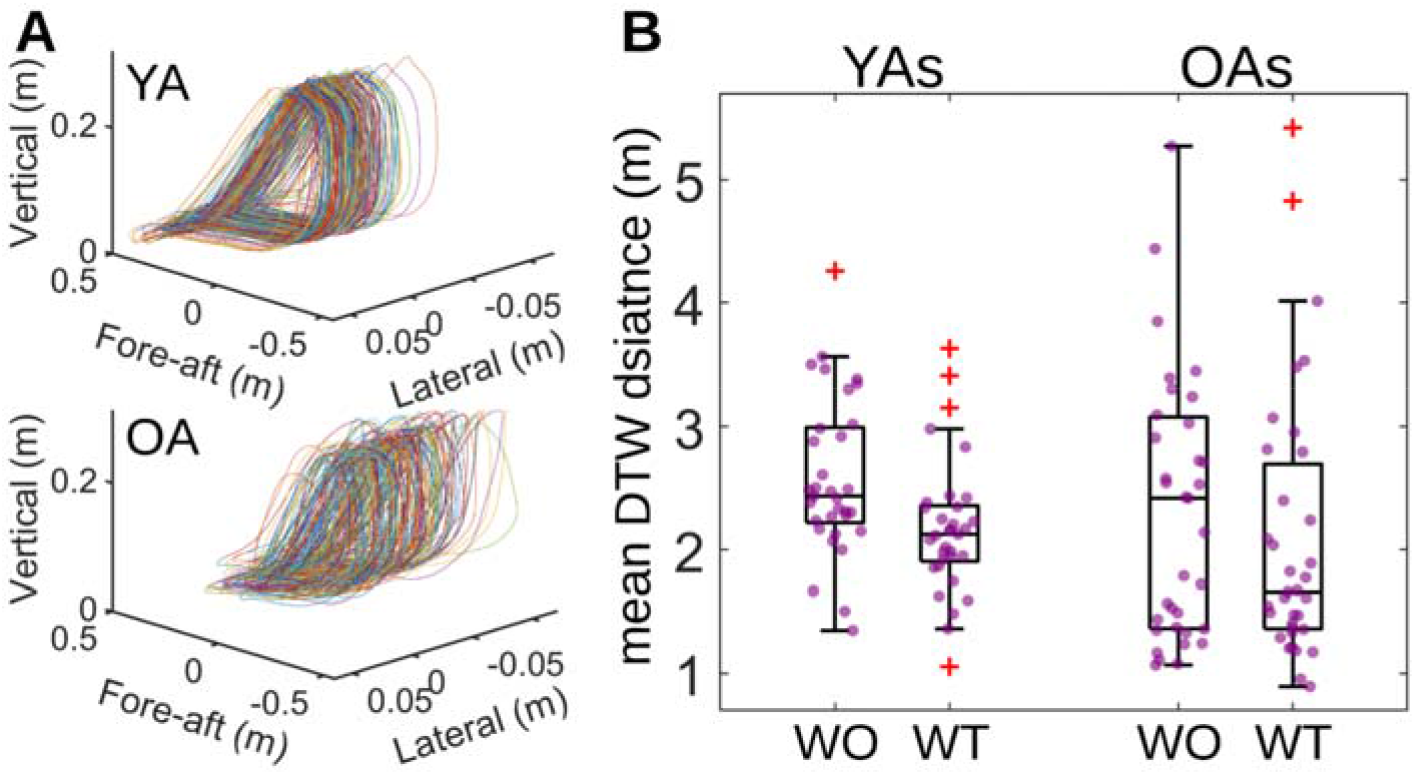
**A**. 3D representations of trajectories of a series of strides for 1 YA and 1 OA, while they concurrently engaged in the Go/NoGo task. Lateral is the dimension of movement right- and-left relative to the motion of the treadmill belt. Vertical is the dimension movement up- and-down relative to the motion of the treadmill belt. Fore-aft is the dimension of movement parallel to the motion of the treadmill belt. Using Dynamic Time Warping (DTW), the variability from one stride to the next was quantified as DTW distance (see Methods) and the mean DTW distance of all stride-to-stride comparisons was extracted per participant. **B**. Mean DTW distance distributions during walking with task (WT) and walking only (WO), in YAs and OAs. Purple dots represent individual participants. Red ‘+’ symbols represent outliers.

Potential associations were tested between WT-minus-WO mean DTW distance and 1) walking-minus-sitting d’ score, controlling for age, as well as 2) age, controlling for walking-minus-sitting d’ score, using partial Spearman rank correlations. Neither of the correlations was found to be significant (Correlation 1: Spearman’s r = -0.10, p = 0.4331; Correlation 2: Spearman’s r = 0.05, p = 0.7060), thereby indicating that the impact of adding Go/NoGo task performance to treadmill walking did not change with age and, also, it was not associated with the impact of adding treadmill walking to Go/NoGo task performance.

### Other Correlations

#### Sex

To investigate whether dual-task-related change in response accuracy could be predicted based on sex, sex was partially correlated with walking-minus-sitting d’ score, controlling for age, using a partial Spearman rank correlation. No significant association between sex and walking-related change in response accuracy was revealed (Spearman’s r = 0.06, p = 0.6205).

#### Cognitive Assessment Score

In older adults, to test whether dual-task-related change in response accuracy could be predicted based on performance in the MoCA cognitive assessment, MoCA score was correlated with walking-minus-sitting d’ score, using a Spearman rank correlation. No significant association between MoCA score and walking-related change in response accuracy was found (Spearman’s r = -0.22, p = 0.1963).

## Discussion

Overall, at the group level, there was a decline in response accuracy while walking that was related to age, consistent with previous studies (1, 17, 21). This relationship was driven by the tendency for a preponderance of the young adults to show response accuracy improvement when walking, whereas most of the older adults showed the opposite pattern, with reductions in walking-related response accuracy. This pattern became evident when walking-related response accuracy distributions were inspected at the single-participant level. In young adults, while there were some participants who declined (DECs; n = 8) or maintained performance (NCs; n = 6) during walking, most improved (IMPs; n = 20). On the other hand, task performance in most older adults declined during walking (DECs; n = 20), while those who improved (IMPs; n = 7) or maintained performance (NCs; n = 10) were fewer, revealing an essentially ‘inverted’ distribution compared to young adults. The young adult distribution was expected based on our prior work (39). However, to our knowledge, response accuracy changes in older adults during walking have not been studied at a single-participant level before. Subsequent analyses aimed to elucidate the finding of ‘paradoxical’ improvement during dual-tasking manifested by these 7 older IMPs, by identifying effects in neurophysiology, gait activity and behavior linked to dual-task-related improvement and aging. Improvement in response accuracy during walking was accompanied by response slowing, suggesting the presence of a speed-accuracy trade-off, as well as by walking-related ERP amplitude modulations during correct rejection trials (i.e. successful response inhibitions). Age was found to correlate with slower treadmill speeds, indicating age-related weakening of ambulatory ability (99-101), and with attenuation in walking-related ERP amplitude modulations during correct rejections.

A clear finding here is that those individuals, regardless of age category, who showed improvement in response inhibition accuracy during walking, were those who showed clear evidence for modulation of their ERP amplitudes across all three tested latency intervals during correct rejections. That is, we observed reductions of the N1 and N2 amplitudes frontocentrally, accompanied by an increase in P3 amplitude lateral-prefrontally (predominantly left), in improving adults during walking (Fig. 5B).

These changes were observed over frontocentral scalp during the N1 and N2 latency intervals, and over predominantly left-lateralized prefrontal scalp during the P3 latency interval (see Fig. 5A). The walking-related ERP amplitude modulations over frontocentral regions during the N2 stage and over prefrontal regions during the P3 stage of correct rejections, which have been previously reported in young IMPs (39), are also apparent in older IMPs (Fig. 5C). Specifically, in young IMPs, reduced walking-related N2 amplitudes frontocentrally have been interpreted as reduced inhibitory conflict during walking, since the source of the N2 has been localized to the anterior cingulate cortex (ACC), which plays a central role in conflict monitoring (54, 55, 58, 59, 102, 103). Also, walking-ERP amplitude modulations over lateral prefrontal regions during the P3 stage have been interpreted as more efficient recruitment of neural resources crucial for top-down behavioral adjustments, which have been localized to the dorsolateral prefrontal cortex, especially in the left hemisphere (39, 51, 102, 104-106).

In contrast to the predictions of CMI, others have proposed that moderate exercise, such as walking, can boost deployment of top-down attentional resources within the lateral prefrontal cortex, possibly mediated by increases in the concentration of catecholamines, serotonin, acetylcholine and cortisol (107-112). More efficient, exercise-induced engagement of these prefrontal resources has been associated with increased arousal levels and improvement in task performance, which likely stems from the adoption of more proactive cognitive strategies to task execution (12, 113-117). Such a shift in the cognitive strategy of IMPs during walking could enhance anticipation of the subsequent ‘NoGo’ trials, thereby explaining why they presumably exhibit walking-related reduction in inhibitory conflict, as indexed by reduced frontocentral-N2 amplitudes (39). The present findings indicate that the neural signatures previously found in young adults who improve during dual-task walking during N2 and P3 (39) also occur in older adults who improve during dual-task walking.

This study additionally revealed a novel neural signature of behavioral improvement during walking, independent of age, specifically the observed walking-related ERP amplitude changes over frontocentral scalp during the N1 stage of correct rejections. The N1 is a negative voltage deflection peaking around 100-200 ms post-stimulus-onset (96-98), and it is thought to reflect sensory gating processes, which involve filtering out irrelevant stimulus information and allocating top-down attentional resources towards enhancing processing of relevant stimulus information (118-121). Neural generation of the N1 has been traced to multiple frontal and parieto-occipital sources, presumably representing the neural centers that top-down attentional control is exerted from and to, respectively (118, 122-124). The reduced walking-related ERP amplitudes over frontocentral scalp regions during N1 latencies manifested by IMPs suggest that, in this behavioral group, walking likely reduced the need for top-down attentional control over task-related sensory and perceptual processes. One possible explanation for IMPs not requiring as much top-down control during the N1 stage is that the neural resources needed at this early processing stage have already been sufficiently activated because of the more efficient frontal resource recruitment exhibited by the same group during the P3 stage of preceding NoGo trials. It is important to highlight that older adults who improved performance while walking also showed these ERP effects during N1 (Fig. 5C). Overall, the neural signatures of improving behavior during N1, N2 and P3 may be due to better management of the inhibitory conflict, which presumably stems from more flexible recalibration of neural processes related to the cognitive component of inhibition, in response to the increase in task demands.

Young adults, regardless of response inhibition accuracy during walking, showed clear modulation of their ERP amplitudes across both tested latency intervals during correct rejections. Specifically, we observed reduction in N2 amplitude lateral-frontally (predominantly left) and parietooccipitally, and reduction in P3 amplitude centroparietally, in young adults during walking. In contrast, such modulations appeared attenuated in older adults, again, regardless walking-related response inhibition accuracy (Fig. 6B).

The topography of these aging-related ERP effects implicates brain regions that are part of the motor inhibition network. Specifically, certain left-lateralized frontal areas, such as the left inferior frontal gyrus and the left lateral orbitofrontal cortex, have been shown to play a key role in implementing top-down inhibitory control over motor responses (53, 65-67, 125). Suppression of motor preparation processes related to pressing the button have been localized to premotor areas contralateral to the hand used to respond during Go trials (65, 66, 126). Since most participants of this study responded with their right hand (59 in total; 30 young and 29 older participants), these neural processes are expected to be reflected left-frontocentrally. Although significant caution must be exercised in inferring intracranial sources from scalp topographic maps, age-related effects when dual-tasking were predominantly found over left frontocentral scalp herein (Fig. 6B). Parietal regions have been long associated with motor attentional control, housing mechanisms related to preparation and redirection of limb movements, and they are particularly activated during tasks that require sequential performance of movements (127-129). During the N2, parietal resources are presumably marshaled to efficiently reorient the motor plan from the prepotent tendency towards motor response execution to the desired motor response inhibition. Motor and midcingulate regions, located centrally, are known to be directly linked to suppression of the prepotent motor response during the P3 processing timeframe (59, 64, 66-69). In the present study, when walking is added to inhibitory task performance, ERP amplitude modulation over these regions in young adults suggests that this age group can flexibly recalibrate neural processes related to the motor component of inhibition. Conversely, attenuation of these modulations in older adults likely indicates that this flexibility is compromised with aging. This interpretation is consistent with previous research reporting age-related functional and structural decline of motor circuitry (130-132). Importantly, these aging-related modulations were present in older adults who improved during walking (Fig. 6C).

Both aging-related and improvement-related neural signatures were manifest over the frontal scalp of improving older adults, during the N2 stage of correct rejections. Comparison of the scalp distribution of the clusters corresponding to each type of signature during the N2 interval shows that the frontocentral behavior-related cluster (red dots on the interval-2 topographical map in Fig. 5C), which has a predominantly midline scalp distribution, and the frontal age-related cluster (red dots on the interval-2 topographical map in Fig. 6C), which has a predominantly left-lateralized scalp distribution, do not overlap in terms of the electrodes that they encompass. This observation further supports the idea that each type of frontal signature during N2 reflects dissociable neural processes underpinned by distinct generators, possibly those discussed in the paragraphs above.

In previous studies, older adults were found to manifest attenuations in walking-related amplitude modulations both for the frontocentral-N2 and for the centroparietal-P3 component (1). In this study, we showed that improving older adults, while they still exhibited such attenuation for the centroparietal-P3 component, they interestingly exhibited walking-related amplitude modulation for the frontocentral-N2 component, which is an effect most commonly encountered in young adults.

One limitation of the present study is that the behavioral effect it targets, namely improvement during walking, was found to have relatively low prevalence in the older adult population (∼19%). As such, despite the relatively large sample size of 71 adults, only 7 older adults were found to show walking-related improvement in the cohort. Future studies could focus on elucidating walking-related improvement in aging by employing screening procedures to specifically identify older adults who improve when dual-task walking, thus increasing the sample size for this group.

## Conclusions

To summarize, aging was found to come with behavioral deterioration in most but not in all adults when combining a Go/NoGo response inhibition task with walking. This study revealed distinct neural signatures of aging and behavioral improvement during dual-task walking. Better response accuracy during walking was found to correlate with slower responses to stimuli, as well as with walking-related ERP amplitude modulations over frontocentral regions during the N1and N2 stages of correct rejections, and over predominantly left-lateralized prefrontal regions during the P3 stage. These neural signatures of behavioral improvement might reflect more flexible recalibration of neural processes related to the cognitive component of inhibition as task demands increase. On the other hand, aging was found to correlate with slower treadmill walking speeds, as well as with attenuation in walking-related ERP amplitude modulations over parietooccipital and predominantly left-lateralized frontal regions during the N2 stage and over centroparietal regions during the P3 stage of correct rejections. These neural signatures likely reflect age-related loss of flexibility in recalibrating neural processes related to the motor component of inhibition with increasing task demands. The smaller percentage of older adults (19%, versus 59% for young adults) who improved performance in the Go/NoGo task during walking exhibited both aging-related neural signatures and neural signatures of behavioral improvement. The coexistence of both types of neural signatures in improving older adults suggests that they likely maintain flexibility in recalibrating the neural processes underpinning the cognitive, but not the motor inhibitory component, when dual-task walking. As such, the neural signatures of behavioral improvement during dual-tasking could potentially serve as markers of ‘super-aging’. Both sets of neural signatures identified in the context of this study hold promise for being translated to clinical populations, such as patients with neurodegenerative diseases, to assess the degree of disease progression, to evaluate treatment outcomes and potentially to identify people, pre-clinically, at high risk for developing aging-related or disease-related cognitive decline.

## Acknowledgements

We would like to thank each of the participants that enrolled in the study.

## Author Contributions

EP: Conceptualization, Data Curation, Formal Analysis, Investigation, Methodology, Software, Visualization, Writing – Original Draft Preparation; JJF: Conceptualization, Funding Acquisition, Methodology, Project Administration, Supervision, Writing – Original Draft Preparation; EPM: Investigation; GK: Investigation; EGF: Conceptualization, Funding Acquisition, Methodology, Project Administration, Supervision, Writing – Original Draft Preparation

## Competing Interests Statement

The authors declare no competing interests.

## Data Availability Statement

Data from this study will be made available through a public repository (e.g. Figshare) upon publication of this paper, and the authors will work with the editorial office during production to incorporate appropriate links. Custom code from this study will be made available on GitHub (https://github.com/CNL-R) upon publication of this paper.

## Funding Disclosure

Partial support for this work came from the University of Rochester’s Del Monte Institute for Neuroscience pilot grant program, funded through the Roberta K. Courtman Trust (EGF).

Recordings were conducted at the Translational Neuroimaging and Neurophysiology Core of the University of Rochester Intellectual and Developmental Disabilities Research Center (UR-IDDRC) which is supported by a center grant from the Eunice Kennedy Shriver National Institute of Child Health and Human Development (P50 HD103536 -JJF). The content is solely the responsibility of the authors and does not necessarily represent the official views of any of the above funders.

## Abbreviations List

CMI: cognitive motor interference
RT: response time
EEG: electroencephalography
ERP: event-related potential
MoBI: mobile brain-body imaging
3D: three-dimensional
DTW: dynamic time warping
IMPs: participants who improved when dual-task walking
NCs: participants whose performance did not change between sitting and walking
DECs: participants who declined when dual-task walking
YAs: young adults
OAs: older adults
CR: correct rejection

## Notes

### Competing Interest Statement

The authors have declared no competing interest.

